# Relationship between the ACTN3, ACE, AGT, BDKRB2 and IL6 genes and the intake of creatine HCl, whey protein and glutamine, with changes in strength and fat percentage, before an undulating strength program in lower limbs in athletes from Valle del Cauca. Colombia

**DOI:** 10.1101/2023.01.14.523685

**Authors:** Gerardo David González Estrada, Efraín Paz, Felipe Sanclemente

## Abstract

Changes in power, strength and muscle mass gain were measured with a group of university athletes (n=11), separating them into two groups, one with supplementation and the other without. supplementation, to determine if the intake of sports supplements had an influence or not on individuals with similar genotypic profiles, or the results of the tests only depended on the predisposition to strength and muscle gain of the ACE, ACTN3, AGT, IL6 and BDKRB2 genes. Genotyping was performed based on PCR, RFLP and polyacrylamide electrophoresis tests. The supplemented group ingested whey protein, creatine HCl, and glutamine. All individuals underwent undulating strength training for four months and jump power tests (SJ, CMJ, and ABA), 1RM, and bioimpedance were performed at three different times.

Changes were obtained in all the athletes, but the group that obtained the greatest gains in all the tests, except the CMJ jump, was the supplemented group and also had a genotypic profile that registered the lowest TGS. In conclusion, we observed significant improvements in individuals with lower TGS and taking sports supplements, surpassing the group that did not take supplements, but had a greater genetic predisposition in strength activities.

## INTRODUCTION

The study of the influence of genes related to physical activity and nutrition has been carried out for approximately 22 years (Ahmad Y. 2020). The analysis of mutations that generate advantages or disadvantages for physical activity and nutrition can be a highly valuable tool for nutritionists and trainers, and must also be taken into account for the development of products in the food industry that maximize sports performance. (Rawson, E. 2018), and the nutritional benefits in certain populations.

The genetic approach for the analysis of physical activity is a valuable tool for decision making about training (Maciejewska-Skrendo 2019). By mixing genetic profiling, field trials, and dietary supplementation, improvement can be achieved in different individual characteristics of recreational or professional athletes (Puthucheary 2011). With the use of models such as the TGS of Williams and Folland 2008, to explain the genotypic variations individual in strength or endurance, we can reveal how genotypes related to physical activity influence different capacities of athletes, such as the maximum amount of oxygen absorbed by the body (VO2 max.), power or maximum strength (Mikami 2014, Saito 2022, Williams 2008) and allows analyzing how these genotype/physical capacity relationships interact, which can be used by physical trainers, coaches and the medical staff, for individual variation in training and improvement of athletes in certain characteristics.

Among the genes that have been related to physical and competitive activity are ACTN3 (Ahmetov I. 2014, Ahmad Y. 2020, Ahmetov I. 2014), ACE (Charbonneau, D. E. 2008, Choudhury, I. 2012), AGT (Gomez-Gallego, F 2009), BDKRB2 (Eynon, N. 2011) and IL6 (Yamin, C. 2008), which are related in their respective order to muscle structure, vasoconstriction, aerobic resistance and muscle repair, being These are among the most studied for these characteristics (Ahmetov, I. 2012).

The objective of the present study is to monitor the changes in a period of four months in a group of amateur athletes, of anaerobic sports (Karate and rugby) and to evaluate the response to tests of power, 1RM and muscle mass gain, between the athletes who were supplemented with Smartnutrition Colombia products and those who were not supplemented, associating this with their genetic profiles, of the ACTN3, ACE, IL6, BDKRB2 and AGT genes.

## METHODOLOGY

### Inclusion criteria

Convenience sampling was carried out among amateur athletes, who compete in the Colombian university cycle, taking the following inclusion criteria, athletes who had continuous training for a year and participated in national category amateur competitions, who did not have sanctions for doping. Personal data, health status and sports performance were taken from them. This project adhered to the protocols required in studies carried out in humans, according to resolution 008430/1993 of the Ministry of health. Prior to taking the sample, each athlete signed an informed consent, where he agreed to participate in this study.

### Sampling and DNA Extraction

The sample was obtained through the buccal scraping technique. Each sample was coded assigning a consecutive number. In the PCR tests, positive controls were used for each allele of each gene and negative controls.

The DNA from the samples was extracted by implementing the protocol used by Quinque et al (2006), for which 2 ml of the mixture of buccal swabs and milliQ water were taken, to which 30 μl of proteinase K and 150 μl of SDS were added. at 10%, leaving this mixture at 57°C for twelve hours. Subsequently, 400 μL of 5M NaCl were added and incubated for 10 minutes on ice. The mixture was distributed in equal volume in 2 mL microtubes and centrifuged for 10 minutes at 13,000 rpm. The supernatant from said process was transferred to a new tube with 800 μL of isopropanol; the tubes were then incubated for 10 minutes at room temperature and subsequently centrifuged at 13,000 rpm for 15 minutes. The supernatants were discarded and the button was washed with 500 μL of 70% ethanol. then it was allowed to dry and dissolved in 40 μL of LowTE.

### PCR amplification of the genes to be evaluated

The genotyping of each one of the loci was carried out by amplification by the PCR technique, from the DNA located in autosomal chromosomes, with the following primers:

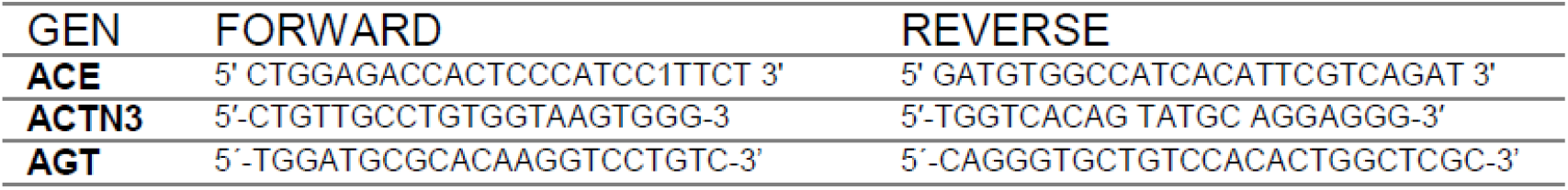

The following amplification protocol was used for all genes, which consists of the following cycles: 95.0 °C, 4:00 min, 35 cycles; 94.0 °C, for 1:00 min; 58.0 °C for 1:00 min; 72.0°C, for 1:00 min. Followed by a final elongation of 72.0 °C, for 10:00 min. The PCR reaction will be carried out in a volume of 25μl, which will contain 3 mM MgCl2, 0.1 mM of primers, 0.1 mM of dNTPs, 1X buffer, 1X Taq polymerase. Enzymatic digestions were performed on the amplified fragments of the ACTN3 and AGT genes with their corresponding restriction enzymes, Dde1, SfnaI.

Electrophoresis was performed in 8% polyacrylamide gels, in TBE1X of the products obtained in the PCR. These electrophoreses were run at 130 volts, and visualized on silver nitrate stained gels.

## PHYSICAL TEST PROTOCOL

Three power measurements were taken in the lower limbs, 1 RM in the lower and upper limbs, and bioimpedance. The data was taken like this, the first one month after starting the undulating strength training protocol, two months after starting the training and four months after finishing the training.

### Power measurement in lower limbs

For alactic anaerobic metabolism, the vertical jump test was used, in which each individual performed a total of 6 jumps, 3 corresponding to the counter movement jump (CMJ), 3 corresponding to the “squat jump” (SJ) and 3 to the jump. abalakov. Each individual performed the tests in the same session, with a rest time of 5 minutes between each one. For the test, measurements were made with the Wheeler Jump photoelectric sensor from the company Wheeler technology applied to sports S.A.S., Colombia.

### Measurement of body composition by bioimpedance

To perform the bioimpedance measurement, a Tanita MC-580 bioimpedance balance was used. The following standards were followed before taking the measurement. Remove all metallic elements from the body (watches, rings, bracelets, earrings, piercings, etc.), do not eat or drink in the 4 hours prior to the bioimpedance test, preferably do not perform bioimpedance in the luteal phase (fluid retention), do not perform strenuous exercise 12 hours before, urinate 30 min. before the test, do not consume alcohol 48 hours before, do not take diuretics 7 days before.

### 1RM measurement

The RM measurement was carried out in the population to be studied, the subjects performed a suitable warm-up before starting to lift the weights. A first series was performed with loads that allowed 6 to 10 repetitions, leaving a minute of rest, before performing the second series which allowed 3 to 5 repetitions, for this second series the increase in weight was between 10 to 20% for the lower body and 5 to 10% for the upper body, after performing this series, he rested for two minutes before performing the third series, which was sought so that only 2 to 3 repetitions could be performed, increasing the weight 10-20% for the lower body and 5-10% for the upper body. They rested for 4 minutes and proceeded to increase the load between 5 to 10% for the upper body and 10 to 20% for the lower body, so that each subject tried to perform a 1RM, if it was possible to move the load, with a appropriate technical gesture, rested again for 4 minutes and increased the load by 5% to try to move it again. When the attempt to do 1 RM fails, rest for 4 minutes and lower the load between 2.5 and 5% for the upper body and 5% for the lower body and again try to move the load.

### Force protocol

Subjects underwent a full body strength training protocol using the undulating methodology. Four weekly sessions were held consisting of exercises for the upper and lower body in each session, alternating daily stimuli in maximum strength, metabolic, hypertrophic and power. The subjects alternated these training sessions with their karate (n=8) and rugby (n=3) practices.

### Supplement intake protocol

The study subjects were divided into two groups of six individuals, one group was given supplements from the Colombian company Smartnutrition, the supplements they consumed were whey protein (a 25-g intake 20 minutes after finishing the strength training), creatine HCl (a 5-g intake 30 minutes before strength training) and glutamine (a 5-g intake in the morning and another 5-g intake at bedtime). Of the group of people who did not supplement, two dropped out due to suffering joint injuries while practicing karate.

### Statistical analysis

A nested experimental design was carried out using as factors the population of elite athletes and that of recreational athletes (control group), race, sex, sport, and specialty of competition.

With the physical tests, the quantitative variables of strength, explosive strength and resistance will be measured, considering in this design the genotypes found for each loci.

The genotypic frequencies of the athletes will be evaluated in compatibility with the Hardy Weinberg equilibrium, the genotypic distribution and allele frequencies between the group of athletes and control will be compared and their significance tested by X^2^ (p <0.05). For the above calculations, the STATISTICA 8.0 software (StatSoft Inc 2007) will be used.

## RESULTS

Table 1 shows the genotypic and allele frequencies of the study groups, the group with supplementation as well as the group that did not supplement, presented a predominance of genotypes and alleles aimed at obtaining good performance in strength. In the last column where the general mean of the group for the TGS test is detailed, the group with supplementation is the one with the lowest value of TGS, which was thus anticipated to provide an observation of the additive effect of the genotypes in the trait of strength for sports performance, people with less predisposition to obtain good results in strength tests were grouped in the group that took supplementation

**Table 1.** Genotypes.

|  | ACE |  |  | ACTN3 |  |  | AGT |  |  | IL6 |  |  | BDKRB2 |  |  | TGS (X) |
| --- | --- | --- | --- | --- | --- | --- | --- | --- | --- | --- | --- | --- | --- | --- | --- | --- |
|  | DD | DI | II | RR | RX | XX | CC | CT | TT | GG | GC | CC | -9-9 | -9+9 | +9+9 |  |
| <u>Supplemented</u> | 2 | 4 | 0 | 4 | 2 | 0 | 3 | 3 | 0 | 4 | 2 | 0 | 0 | 6 | 0 | 71 |
| <u>Not Supplemented</u> | 3 | 1 | 0 | 2 | 2 | 0 | 3 | 1 | 0 | 3 | 1 | 0 | 1 | 3 | 0 | 80 |

Table 2 details the changes in power with the SJ, CMJ, and ABA jumps. In maximum strength with the 1RM tests in the squat, bench press, shoulder press and deadlift. In addition to measuring the change in muscle mass by bioimpedance test. For the SJ, CMJ and ABA jumps, the greatest gains in height are observed by individuals of the group with supplementation like this, the individuals SP1, SP2 and SP3 for the SJ. The individual SP1 for the CMJ and the individuals, SP1, SP2, SP5 and SP6 for the ABA. In the 1RM tests, the best results occurred in the group of supplemented individuals, without any being below the lowest result obtained in the group without supplementation. In the results of muscle gain, the supplemented individuals presented mostly (four out of six) positive gains in muscle mass and only two presented mass loss. However, in the group without supplementation the results were mostly negative (three out of four). It should be noted that the individual supplemented with SP1, despite having the lowest TGS of all the participants, managed to outperform all the individuals in both groups, in most of the tests, and was the one who obtained the greatest gain in kg of muscle mass. of all the participants.

**Table 2.** Gain in cm and kg in the tests performed.

| GROUPS | INDIVIDUAL | GAIN IN JUMP HEIGHT IN<br>Cm |  |  | WEIGHT LIFTING GAIN final kg - initial<br>kg |  |  |  | MUSCLE<br>MASS<br>GAIN Kg | TGS |
| --- | --- | --- | --- | --- | --- | --- | --- | --- | --- | --- |
|  |  | SJ | CMJ | ABA | <u>Squat</u> | <u>bench</u><br><u>press</u> | <u>deadlift</u> | <u>Shoulder</u><br><u>press</u> |  |  |
| **Supplemented** | SP1 | 11,5 | 10,6 | 14,3 | 32 | 25 | 26 | 10 | 3,2 | 50 |
|  | SP2 | 7,4 | 3,2 | 9 | 12 | 29 | 27 | 15 | 1,6 | 70 |
|  | SP3 | 6,4 | -0,5 | 6,2 | 14 | 8 | 18 | 7 | -0,1 | 90 |
|  | SP4 | 1,2 | 3,9 | 1,6 | 32 | 18 | 42 | 8 | 0,6 | 60 |
|  | SP5 | 0,5 | 3,6 | 6,9 | 26 | 11 | 27 | 8 | -1 | 80 |
|  | SP6 | 0,7 | 5,2 | 14,4 | 38 | 25 | 19 | 19 | 1,7 | 80 |
| **not supplemented** | SP7 | 7,6 | 10,5 | 2,6 | 20 | 11 | 19 | 7 | -0,3 | 90 |
|  | SP8 | 0,4 | 5,3 | 5,6 | 6 | 8 | -10 | 2 | -3,3 | 90 |
|  | SP9 | 3,9 | 10,5 | 6,9 | 35 | 6 | 18 | 7 | -2,8 | 80 |
|  | SP10 | 3,3 | 2,3 | 5,4 | 13 | 7 | 7 | 2 | 0,6 | 60 |

Table 3 compares the means of the group of supplemented and non-supplemented people, in the different tests, it can be seen that the means of the group of supplemented people exceed the nonsupplemented group in all tests, except in the test of CMJ jump. Significant differences are observed in the tests related to the measurement of the 1RM, in the rest of the tests no significant differences are reported.

**Table 3.**
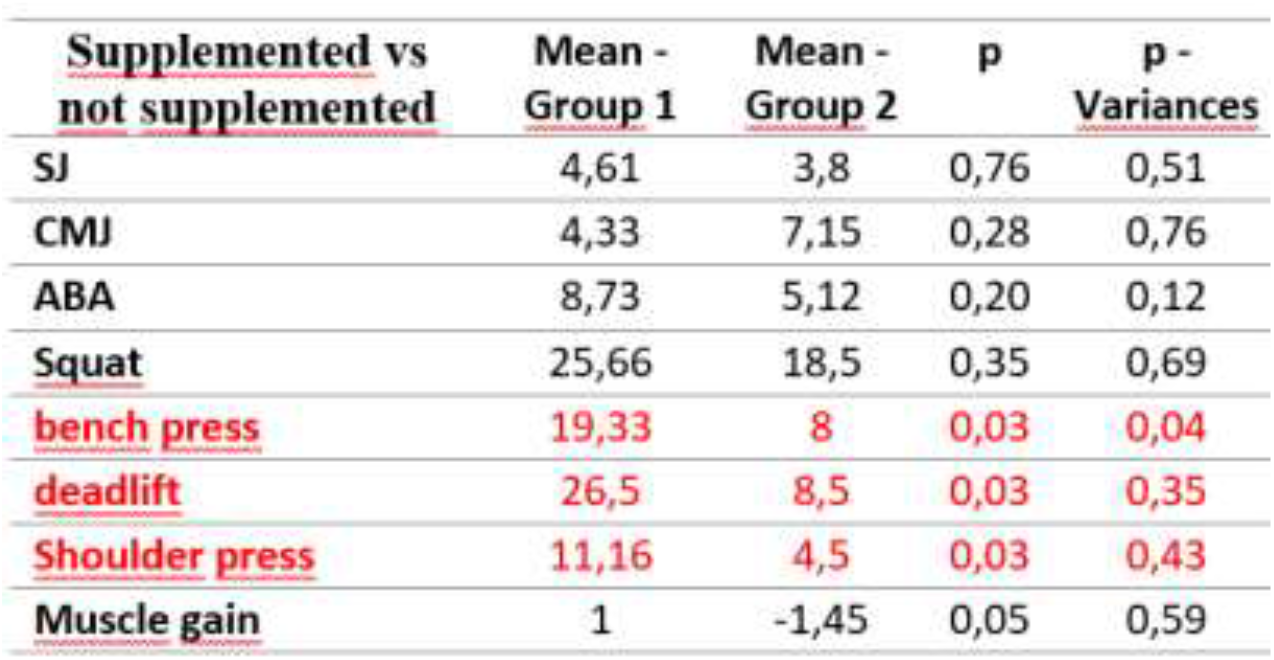
Comparison between the group of supplemented and non-supplemented individuals in the different strength and muscle gain tests.

| **<u>Supplemented vs</u>**<br>**<u>not supplemented</u>** | **<u>Mean -</u>**<br>**<u>Group 1</u>** | **<u>Mean -</u>**<br>**<u>Group 2</u>** | **<u>p</u>** | **<u>p -</u>**<br>**<u>Variances</u>** |
| --- | --- | --- | --- | --- |
| **SJ** | 4,61 | 3,8 | 0,76 | 0,51 |
| **CMJ** | 4,33 | 7,15 | 0,28 | 0,76 |
| **ABA** | 8,73 | 5,12 | 0,20 | 0,12 |
| **Squat** | 25,66 | 18,5 | 0,35 | 0,69 |
| **<u>bench press</u>** | **19,33** | **8** | **0,03** | **0,04** |
| **<u>deadlift</u>** | **26,5** | **8,5** | **0,03** | **0,35** |
| **<u>Shoulder press</u>** | **11,16** | **4,5** | **0,03** | **0,43** |
| **Muscle gain** | 1 | -1,45 | 0,05 | 0,59 |

## DISCUSSION

Changes were observed in all the study participants, when comparing the initial and final time, but the group that took the supplements, presented mostly an increase in the measurements of the SJ, CMJ and ABA jumps, which the majority of individuals from the group without supplementation, despite this at a statistical level did not present significant differences, this probably due to the number of people studied, but these gains in cm, in competition, do represent a difference that would lead to obtaining or not an ideal sporting result.

It is noteworthy that despite the fact that the genetic profiles of the majority predispose them to obtain good results in strength tests (TGS>70), (Cieszczyk 2011, Gineviciene 2016, MacIejewska-Skrendo 2019), the people in the group with supplementation had profiles with lower TGS, taking into account that the higher the measured TGS, the more predisposition to strength the individual presents (Buxens 2010), despite this the SP1 individual, who had the lowest (TGS=50), but supplemented by ingesting whey protein, creatine HCl and glutamine, surpassed the individual of the same sex (SP7) in most jump tests and 1RM but with a much higher TGS (TGS=90). Showing how an epigenetic influence, related to strength training accompanied by adequate supplementation, can maximize the genetic influence to a great extent and results can be achieved that surpass individuals with a greater genetic predisposition in strength tests, but with influences on not so adequate intake (Maughan 2018). In general, the group with supplementation outperformed the group without supplementation except in the CMJ jump test as shown in table 2.

In the bioimpedance test, the changes in muscle mass gain were greater in the group that consumed whey protein, creatine HCl, and glutamine, showing in three of the four individuals in the group without supplementation, a loss in muscle mass, despite to maintain a protein intake in your daily diet. This shows how in this study, the consumption of ergogenic aids and protein was of vital importance, for the improvement in the gain of muscle mass (Valenzuela 2018). What is very important for sports practitioners at a professional or amateur level, as well as for people who practice it for health or fitness.

Despite the fact that the number of study participants is low, which limits the power of statistical analysis, it was possible to observe how there were marked differences in the means between the participants, in seven of the eight tests, and significant differences in all the 1RM measurement tests, in addition the gain in muscle mass measured by bioimpedance was also higher in people with supplementation (Schoenfeld 2018).

All these differences occur in individuals with genotypic profiles aimed at having good results in strength tests, making the group quite homogeneous, since the ACTN3 and ACE genes, which have been one of the most studied in sport genetics, show in previous analyzes, that the DD and DI genotypes of the ACE gene and the RR and RX genotypes of the ACTN3 gene are those that generate the best predisposition in strength activities (Berman 2010), and precisely these genotypes are those presented by the individuals of the two groups, which that gave a general orientation of the participants to have a good performance in strength activities. In addition, the IL6 gene that has been associated with orientation to strength activities and muscle recovery (Yamin, C. 2008), does not present in any of the participants the TT genotype that has been registered as unfavorable for the two aforementioned qualities. The genotypes that predispose to have better performance in aerobic activities such as the II of the ACE gene, the TT of the AGT gene and the XX of the ACTN3 gene did not appear in any individual (de Moor 2007, Grenda 2014), and the genotype −9 −9 of the BDKRB2 gene (Grenda 2014) only occurred in a single individual. Therefore, the influence that was given, by supplementation and strength training, influenced in a general way individuals with a predisposition to obtain good results in strength tests.

In conclusion, the study allowed us to observe how the influence of ergogenic aids and protein (Stokes 2018) maximize the genetic predisposition of individuals, leading them to obtain better results in strength tests (Luzi 2012), surpassing individuals with the same or better genotypes (TGS) related to strength activities, such as those of the non-supplemented group, in addition to the fact that muscle mass gain is also greatly benefited in supplemented people (Deldique 2020), despite the fact that those not supplemented present better or equal genotypes oriented towards hypertrophy, such as the DD of the ACE gene, the GG of the IL6 gene and the RR of the ACTN3 gene. It is recommended for future studies to increase the number of individuals, and thus give greater power to the statistical analysis.

## ACKNOWLEDGMENTS

To the company Smart Nutrition Colombia for collaborating with its sports supplement products to carry out this study.

## Notes

### Competing Interest Statement

The authors have declared no competing interest.

